# Risk factors and spatiotemporal analysis of classical swine fever in Ecuador

**DOI:** 10.1101/2022.09.02.506027

**Authors:** Alfredo Acosta, Klaas Dietze, Oswaldo Baquero, Germana Vizzotto Osowski, Christian Imbacuan, Lidia Alexandra Burbano, Fernando Ferreira, Klaus Depner

## Abstract

Classical swine fever (CSF) is one of the most important re-emergent swine diseases worldwide. Despite concerted control efforts in the Andean countries, the disease remains endemic in several areas, limiting production and trade opportunities. In this study, we aim to determine herd-level risk factors and spatiotemporal implications associated with CSF. We analysed passive surveillance datasets and vaccination programmes from 2014 to 2020; Then, structured a herd-level case-control study using a multivariable logistic model containing 339 cases, and a spatiotemporal Bayesian model, considering 115 thousand premises, 2.3 million annual vaccine doses and a population of 1.6 million pigs distributed in 1,006 parishes. Our results showed that the risk factors that increased the odds of CSF occurrence were swill feeding (OR 9.28), time of notification (OR 2.18), animal entry in the last 30 days (OR 2.08), lack of CSF vaccination (OR 1.88), age of animals between 3-6 months (OR 1.58) and being in the coastal region (OR 1.87). Spatiotemporal models showed that the vaccination campaign reduced the risk by 33% while temperature increased the risk by 17%. The calculated priority index aims to facilitate the intervention process that should be focused on a couple of provinces, mainly in Morona Santiago and Los Rios as well as in specific parishes around the country. Our findings provide insight and understanding of the risk factors associated with CSF in Ecuador, which stands for the Andean region; even though the results are specific for the implementation of risk-based surveillance for CSF, data and methods could be valuable for the prevention and control of diseases such as African swine fever, or porcine reproductive and respiratory syndrome. In conclusion, the results highlight the complexity of the CSF control programme, the need to inform decision-makers, involve stakeholders and implement better strategies to update continental health policies to eradicate swine diseases.

## 2. Introduction

With the growing demand for animal protein in the Andean region, especially pork, diseases such as classical swine fever (CSF) are gaining importance as they are limiting local production and potential export opportunities for affected countries. The per capita consumption of pork has increased in Ecuador from 7.3 kg in 2010 to 10 kg in 2016 (www.aspe.org.ec). The overall importance of pig production is related not only to meat consumption but also to cultural traditions. In Andean communities, pigs play a central role as a source of protein, festivities and savings (1).

Classical swine fever is considered the most relevant re-emerging viral disease of pigs and is caused by a virus belonging to the *Pestivirus* genus within the family *Flaviviridae* (2). The only natural reservoirs are members of the *Suidae* family (domestic and wild pigs). Clinical signs are variable and depend on the viral strain, host immune response, age, general health status and concomitant infections (3).

The presentations of the disease include acute, chronic and persistent forms according to their duration rather than their different acute, chronic and persistent manifestations (4,5). Transmission occurs mainly by direct contact between infected and susceptible animals via the oronasal route but also indirectly through people, clothes, vehicles, equipment, and ingestion of contaminated and undercooked meat as part of swill feeding (6). Outbreaks of CSF usually have dramatic consequences when control measures are implemented. These include long quarantine periods, movement restrictions, emergency vaccination, culling of the pig population and major impacts on animal welfare (7,8). For instance, the 1998 epidemic in the Netherlands, had an estimated cost of 2.3 billion US dollars and the destruction of 10 million pigs (9). Countries with endemic status are banned from exporting pigs and their products, therefore the impact of the disease on the economy and public health worldwide is enormous. In Ecuador, the economic impact caused annually by CSF was estimated by the NVS to be 6 million US dollars, considering only animal mortality. As many of the involved people are from households with low income, the impact of CSF on particular poor people becomes evident.

In South America, the disease is considered endemic in Guyana, Suriname, the North and Northeast regions of Brazil and the Andean Community, and these regions struggle to implement successful control programs (10,11).

The national CSF eradication Project in Ecuador was established in 2012 with significant improvements against CSF by the National Veterinary Service (NVS). The first national vaccination strategy was gradually initiated in 2014 with a locally produced lapinised Chinese strain vaccine (12). As a result, the highest historical coverage was achieved in 2019 (2.7 million doses) due to a compulsory vaccination campaign, government subsidies and coordination with stakeholders (commercial and industrial producers’ associations). However, the field response and data analysis capacity of the veterinary service is limited, and in 2020, the disease is still present (https://wahis.oie.int). In this regard, the NVS plans to enhance their analysis capacity and apply riskbased surveillance. One of the main challenges in applying risk-based surveillance is to find the factors associated with the occurrence of a given disease; in the case of CSF, the disease and associated risk factors have been well studied for developed countries (13–18). However, in developing countries, where due to very particular production systems, the risk factors may be different, these are yet to be understood. Recently, for some countries in South America such as Colombia (19), Brazil (20), and Peru (21), this issue has been addressed.

Despite the importance and need for local CSF risk factors, little is still known about them in Ecuador, generating a lack of knowledge to inform control measures and public policy. This is the first time that official data from the NVS has been analysed to determine risk factors for the period from 2014 to 2020.

The objectives of this study were to determine the risk factors associated with the occurrence of CSF and to analyse the spatiotemporal implications in order to identify the regions most at risk.

## 3. Materials and methods

### 3.1 Datasets

Data were collected by the NVS from January 2014 to November 2020 in mainland Ecuador, excluding the Galapagos Islands, as it is a recognised CSF free zone (2). The information was stored in two databases: (1) Ecuador’s animal health information system (SIZSE) created to record paper questionnaires for notifiable diseases, suspicious events and laboratory results from passive surveillance since 2014 (https://sistemas.agrocalidad.gob.ec/sizse/); and (2) the Unified information manager (GUIA) developed by the NVS to manage cadastre, mass vaccination campaign against CSF, and movements since 2016 (https://guia.agrocalidad.gob.ec/agrodb/ingreso.php). Shapefiles of administrative units of Ecuador were downloaded from the Institute of Statistics and Census (INEC) (https://www.ecuadorencifras.gob.ec/division-politico-administrativa/).

All raw data were then imported and processed with R version 4.2.1 (https://CRAN.R-project.org/).

### 3.2 Factors influencing the risk (variables captured by the surveillance system)

The retrospective analysis used the variables collected historically by the surveillance system. The databases were merged using the individual identification of the owner of the premises.

The variables included in the analysis were selected based on published literature, biological plausibility and considering their association with the occurrence of classical swine fever. (14,22,23). Subsequently, they were organised considering risk characteristics according to the RiskSur Surveillance design framework (www.fp7-risksur.eu) (24) and classified into the population level, herd level and animal level.

Using the variables included in the passive surveillance system, a case-control study was structured to identify factors associated with CSF occurrence. Herds of pigs with a positive laboratory test for CSF were classified as case herds, and those with a negative result were classified as controls (Figure 1).

**Figure 1.**
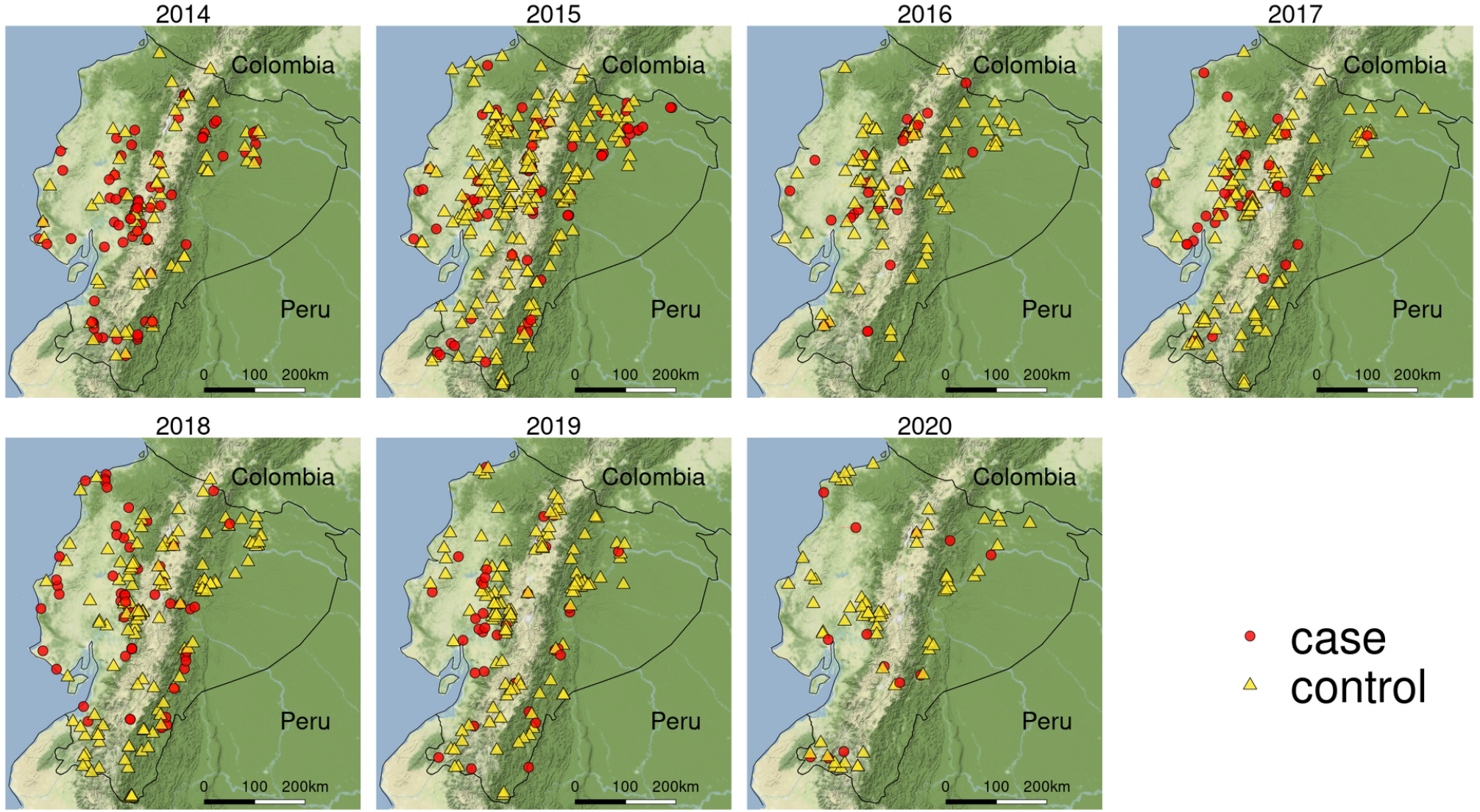
Spatial representation of the study area and location of cases and controls of CSF in Ecuador study period 2014–2020.

### 3.3 Questionnaire

The surveillance system used a questionnaire designed to obtain information on veterinary health events, including the demographic data of the premises, geographic location, chronology (dates of notification and follow-up), animal species, vaccination declaration, clinical signs, presumptive syndrome, collection of material, characteristics of samples, laboratory test results, animal population, animal movement and probable origin of the disease.

The information was collected by trained NVS veterinarians following the data protection procedures of Ecuadorian authorities. The information recorded throughout the country was continually monitored by the national surveillance team (headquarters), who checked the data for completeness and errors.

Laboratory testing was performed at the National Reference Laboratory (headquarters in *Quito*). Virus detection was carried out by a commercially available antigen ELISA (PrioCheck® CSFV) based on the double antibody sandwich (DAS) principle with a sensitivity of 97% and a specificity of 99% (25) and by qRT-PCR using Roche® reagents (26) with a sensitivity and specificity of ≥ 95%.

### 3.4 Multivariable logistic analysis

All analyses were performed at the herd level and stratified according to CSF status (case or control). Variables were organised by type, continuous variables were transformed into dummies, setting their levels according to biological or legal cut-off points. Descriptive statistics assessed the distribution of cases and controls. The dependent variable was the binary variable infected with CSF. The evaluation of individual variables of the Ecuadorian surveillance system was based on the association of each explanatory variable with the binary farm-level outcome, using univariate logistic regression (27). We avoided case-control matching due to the potential of creating selection bias, losing precision, statistical power and not having a prior local analysis of strong well-measured confounding variables (28).

A multivariate logistic regression model (Eq. 1) was used to assess the association of explanatory variables with the outcome formulated as follows:

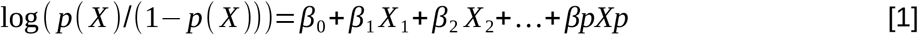

where X_j_: is the jth predictor variable and β_j_ is the coefficient estimate for the jth predictor variable. The final model selection used a manual forward stepwise approach (29). We included each variable in descending order of statistical significance in the univariate models. Statistically significant variables (chi-square association test < 0.05) were kept in the final model. For each insertion of new variables, we observed the changes in the odds ratio (OR) and the significance of each beta β_i_ (Wald test) assessing them at each step. Variables were used only if their completeness was ≥ 0.85. Confounding was assessed using causal diagrams (30,31). Collinearity was analysed using variance inflation factors analysis (32). The goodness of fit of the final model was measured using the conditional R_2_ (33) and the Hosmer-Lemeshow goodness of fit test (GOF) (p > 0.05) (34).

### 3.5 Spatiotemporal Bayesian analysis

The analysis used the population and cases restricted to 2017-2020 due to the lack of cadastral and vaccination information prior to the implementation of the increased official vaccination in 2017. Data were organised to contain the aggregated annual population over each parish (1040) using time-series missing value imputation (35) for areas without information. Variables were centred and scaled by dividing the centred value by the standard deviation. The variables used to fit the model were the number of CSF vaccine doses applied per km^2^, average temperature (C) and average precipitation (mm), constructing several models. Temperature and precipitation were extracted from (https://worldclim.com/) at a spatial resolution of 2.5 arc-minutes (∼5 Km^2^).

Parish vaccination coverage was adjusted considering the population and the doses applied against CSF, considering 1.55 as the average number of doses a pig receives in a calendar year, according to the average lifespan from birth to slaughter (234 days) (36). We used penalised priority priors model complexity, specified by the divergence between a flexible model and a baseline model; To define the spatial random effect, a neighbourhood matrix from the polygon list was needed, based on regions (parishes), that share two or more boundary points. The spatiotemporal model uses the disease count Y_ij_ observed in area i and time period j, modelled as:

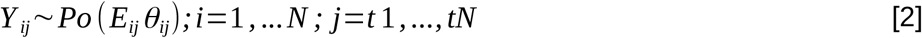

where *E*_*ij*_ is the expected number of cases and *θ*_*ij*_ is the relative risk, both in the given area (*i*) and time period (*t*) (Equation 2). Three sets of components for log (*θ*_*ij*_) were considered:

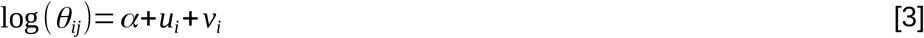

where alpha represents an overall risk in the study region, *u*_i_ is the correlated heterogeneity that models the spatial dependence between the relative risks, and *v*_i_ is the unstructured exchangeable component that models uncorrelated noise (Equation 3).

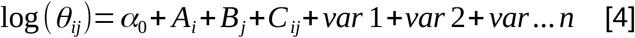

where A_i_ represents the spatial group, B_j_ is the temporal group, and C_ij_ is the space-time interaction group (A_i_ = u_i_ + v_i_) using the most popular model to spatial disease known as Besag-York-Mollié (BYM) (37), where the clustering component *u*_i_ is modelled with the conditional autoregressive distribution (CAR) (38), smoothing the data when two areas share a common boundary given by the neighbourhood matrix (B_j_ = βt_j_). Using an independent and identically distributed Gaussian random effect (iid). (C_ij_ = δ_i_t_j_), where u_i_ + v_i_ is an area random effect, βt_j_ is a linear trend term in time t_j_, and δ_i_t_j_ is an interaction random effect between area and time (Equation 4) (39).

To evaluate the models, we used the deviance information criterion (DIC) and the posterior predictive p value. To suggest parish in priority of care, we used the *priority index* (PI) which is a risk-based percentage scale that ranks the units of analysis, given by the fitted effects weighted by their probability and a cut-off value (40). The models were implemented using the integrated nested laplace approximation (INLA) (41). We used choropleth maps to represent the spatiotemporal distribution of the population, observed cases, expected cases, infection risk (relative risk) and priority index.

All analyses were run in R V4.2.0.

## 4. Results

### 4.1 Descriptive analysis of the variables influencing the risk

The full dataset contained 63 variables, most of which were used for administrative purposes. Fifteen variables selected for the univariate analysis consisted of 6 dichotomous, 6 nominal and 3 continuous variables. They were then classified into the population level (n=4), herd level (n=9) and animal level (n=2). Notification time was transformed into a dummy variable considering a cut point of 7 days (one week). The age of the animals was considered cut-off points of 2 and 6 months due to the official CSF vaccination recommendation: first dose applied after 45 days of age and revaccination at 180 days of age (Table 1).

**Table 1.** Description of variables influencing risk and their levels. Data available in Ecuador’s CSF surveillance system from 2014 to 2020, levels grouped by risk characterisation.

| Factors influencing risk | Description of variables captured by the surveillance system | Category |
| --- | --- | --- |
| <sup>†</sup>Control program | Active national control program (vaccination and mobilisation control) on the movement of interview. | Dichotomic <sup>¶</sup> |
| <sup>†</sup>Natural region | Natural region of the premise according to their location and administrative provincial division. | Amazon, coastal, highlands |
| <sup>†</sup>Network community | Community to which the premise belongs according to its parish location (42). | 5 communities |
| <sup>†</sup>Year | Year of the event. | 2014–2020 |
| <sup>‡</sup>Animal entry | Reception of pigs within 30 days of onset of clinical signs. | Dichotomic <sup>¶</sup> |
| <sup>‡</sup>Notification time | Number of days from onset of clinical signs to notification to the NVS. | 0–7, >7 |
| <sup>‡</sup>Other species | Existence of species other than swine in the premise. | Dichotomic |
| <sup>‡</sup>Size of the farm | Number of pigs on the premises. | 1–25, 26–189, >190, |
| <sup>‡</sup>Swill feeding | Evidence of feeding pigs with swill feed, home-made leftovers. | Dichotomic <sup>¶</sup> |
| <sup>‡</sup>Type of farm | Classification of premises according to production category. | Backyard, Family, Commercial, Industrial |
| <sup>‡</sup>CSF vaccination declaration | Owner's declaration of vaccination against CSF in its premise. | Dichotomic <sup>¶</sup> |
| <sup>‡</sup>CSF vaccination record | Official record of vaccination against CSF within the last 180 days. | Dichotomic <sup>¶</sup> |
| <sup>‡</sup>Who makes the notification | Person who contacted the NVS to make the notification. | Owner, NVS, Sensor |
| <sup>§</sup>Age | Age in months of sampled animals. | 1–2, 3–6, >=7 |
| <sup>§</sup>Breed | Breed of the animals on the premise. | Landrace (white), Indigenous (black). |

Levels of risk characterisation: <sup>†</sup> Population level, <sup>‡</sup> Herd level, <sup>§</sup> Animal level. <sup>¶</sup> Dichotomic: 0=No, 1=Yes.

The average time for official notification to the NVS was more than one week (9.3 days) for cases and one week (7.0 days) for controls. The median farm size was similar for cases (13.5 pigs) and controls. The mean age of pigs was similar for cases and controls (∼5 months) (Table 2).

**Table 2.**
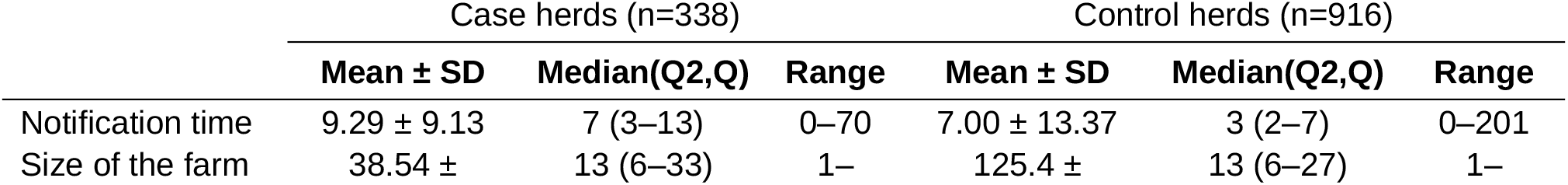
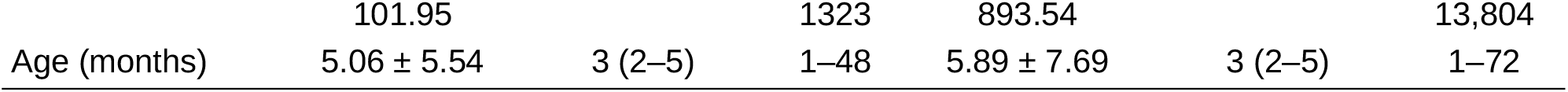
Descriptive measures of continuous variables from the 2014–2020 CSF risk factor analysis in Ecuador.

Farmer vaccination declaration was higher for case herds (71%) than for control herds (6%). Recording of vaccination (based on official records) was lower (46%) for cases than for controls (60%). Both cases (96%) and controls (78%) had a high percentage of swill feed use. The entry of animals within the last 30 days was recorded in 39% of cases and 22% of controls. Only 5% of the cases and 4% of controls have other species on the property (Table 3).

**Table 3.**
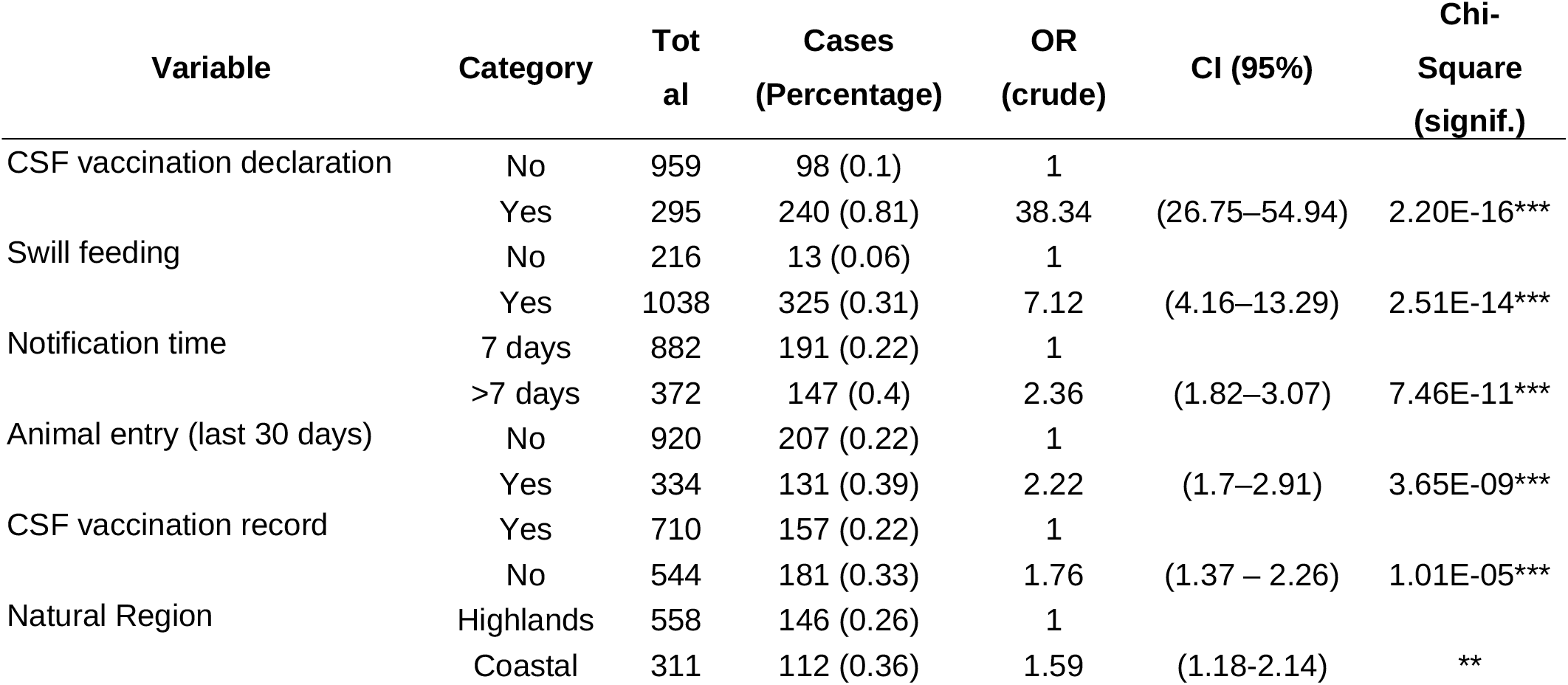
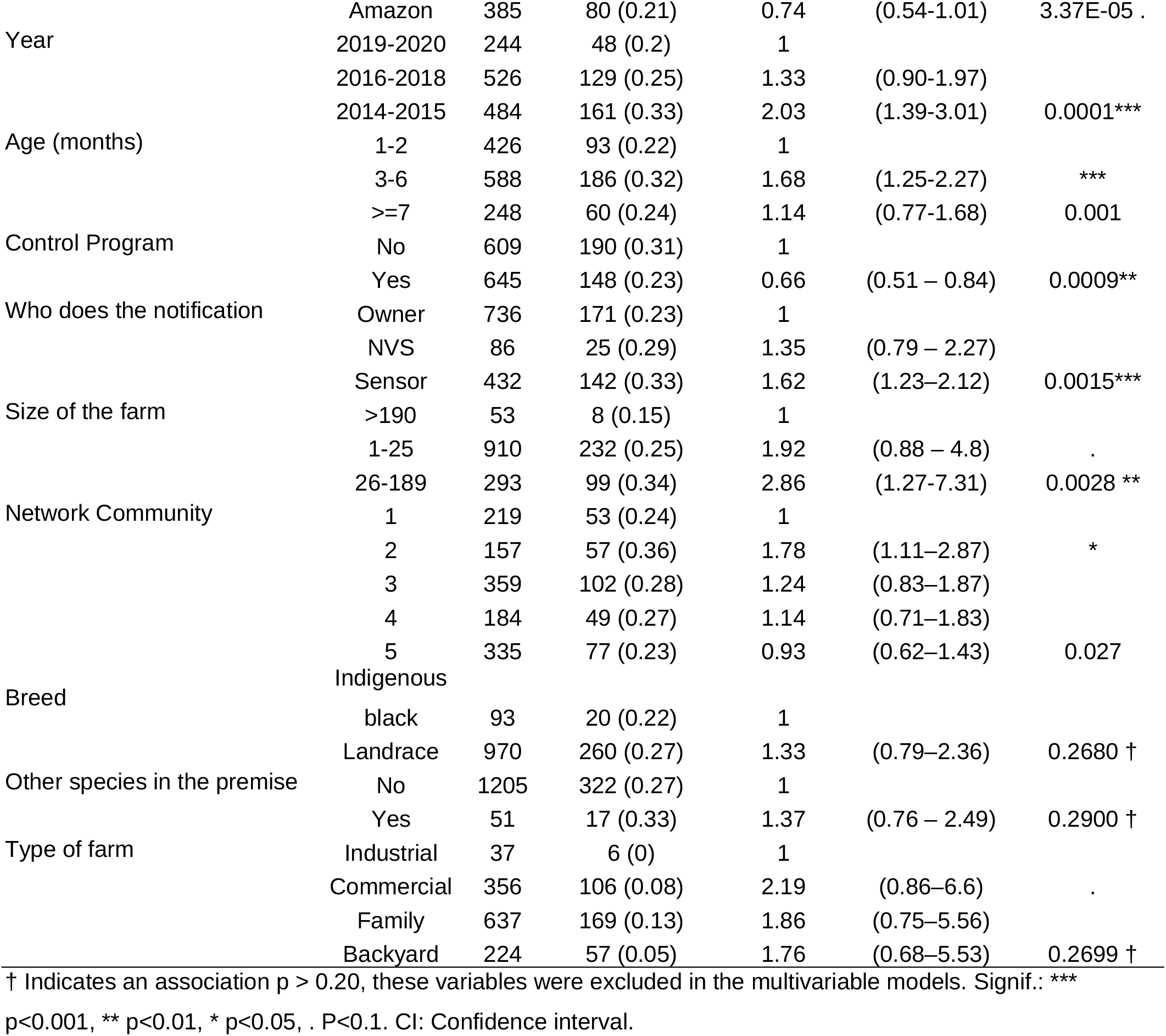
Results of univariable logistic regression analyses, to assess associations of CSF during 2014–2020 in Ecuadorian swine herds. Variables are ordered by their level of significance.

Historical case presentation decreased over the years, with the highest proportion of cases (48%) occurring between 2014 and 2015 and the lowest (14%) occurring between 2019 and 2020. The proportion of cases (43%) and controls (45%) was highest in the highlands. More than half of the case notifications (51%) were reported by the owner, followed by health sensors (42%), which are volunteers selected by the NVS to enhance the surveillance system directly from the community across the country. There was a higher proportion of cases (30%) in the third community network, located in the centre of the country. The highest proportion of cases was in commercial production (31%) (Table 3).

### 4.2 Description of cases and controls

The surveillance database contained 1,254 questionnaires, 338 of which were confirmed CSF cases. The farm categories were 50.79% family (637), followed by 28.39% commercial (356), 17.98% backyard (224) and 0.03% industrial (37); the distribution of cases and controls over time is illustrated in Figure 2; The highest case presentation corresponded to October 2015 with 14 monthly cases, followed by March 2014 with 13 cases, and the lowest corresponded to 2020 with ≤ 2 monthly cases.

**Figure 2.**
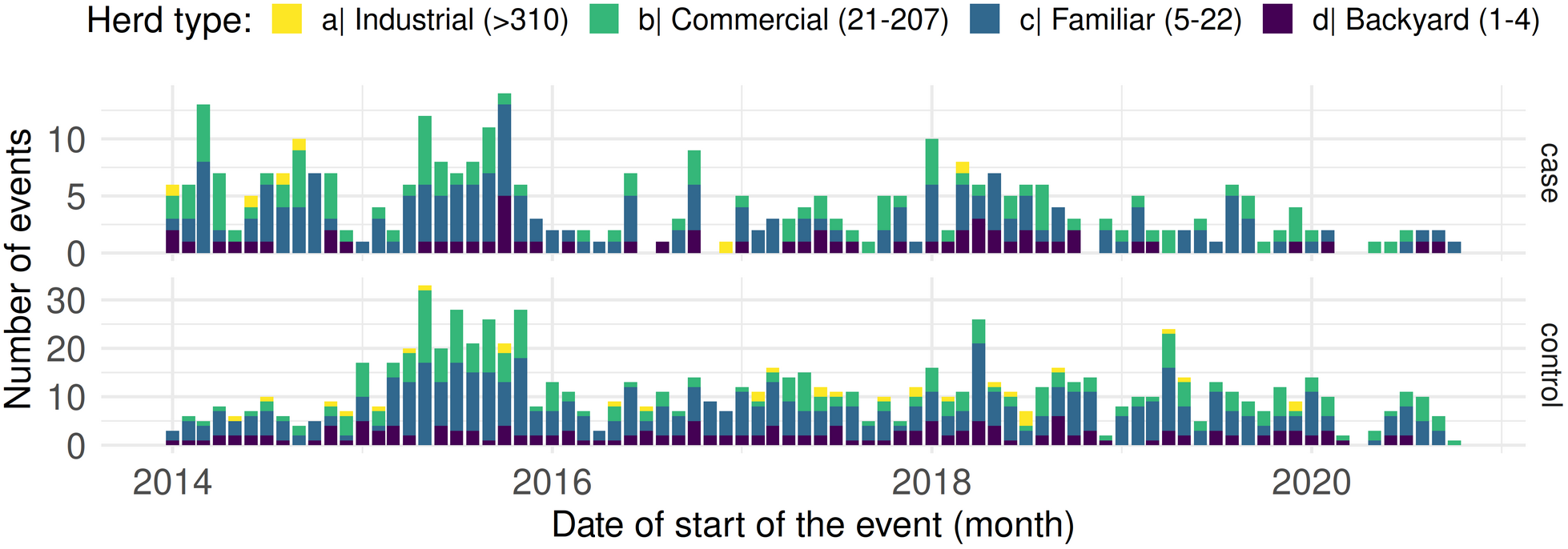
Distribution of events (cases and controls) reported between 2014 and 2020 in Ecuador; bars represent monthly counts, height and colours of bars according to the type of herd (different scales on the y-axis).

Distributed by months, the mean number of cases was 4.39 ± 3.09; with controls, the mean was 11.31 ± 6.38 There was a tendency to increase twice a year around April and October (Figure 3).

**Figure 3.**
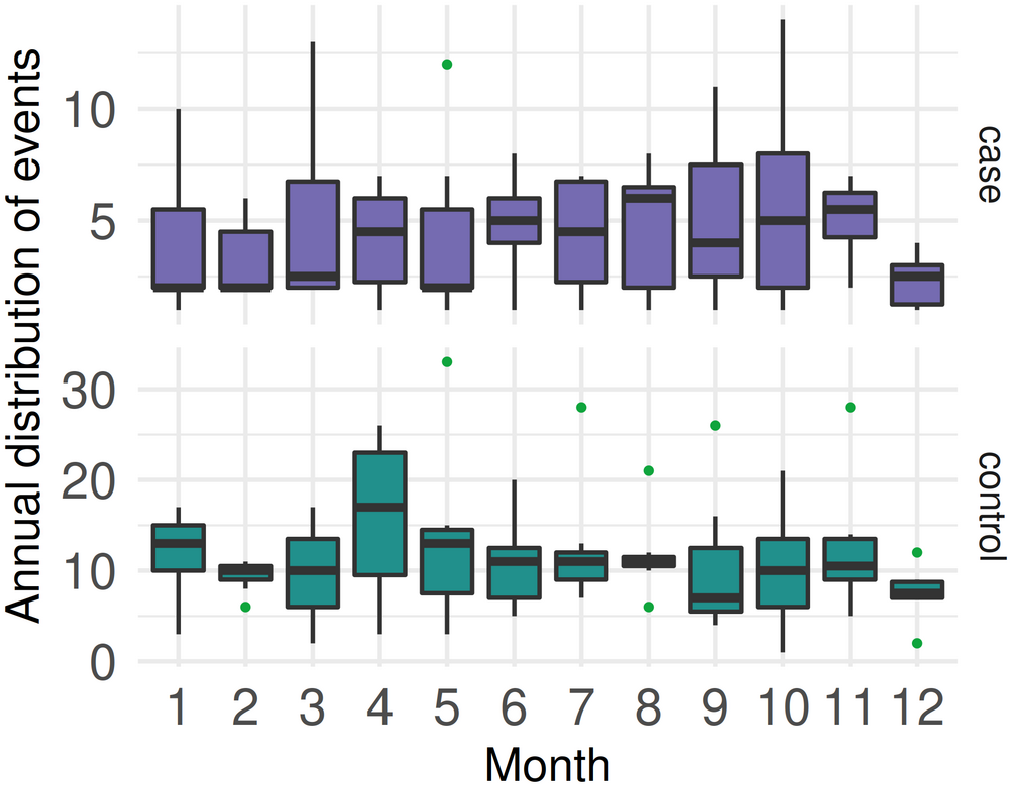
Boxplots of monthly distribution of CSF cases and controls from 2014 to 2020 in Ecuador (different scales on the y-axis).

### 4.3 Multivariable logistic analysis

According to the univariable models, twelve out of 15 assessed variables were associated with CSF (p<0.20). We found a paradoxical fit (Type III error) opposite of the true effect (42), produced by CSF vaccination declaration; giving an incorrect direction of association and increasing the odds when the farmer declares vaccination (38.67 OR), instead of the expected protective effect conferred by the vaccine. The natural region showed a higher risk to the coastal region. The univariable analysis is presented in Table 3.

During the stepwise forward selection of variables we evaluated 8 models; thus, variables that were not statistically significant (p>0.05), or had an incorrect direction of association (self-declaration of vaccination) were excluded. Variables that showed a strong association (p<0.0001) in the univariable model, maintained their individual and model significance when adjusted in the final multivariable model (Table 4).

**Table 4.** Multivariable logistic regression model assessing the associations of variables with the odds of CSF between 2014 and 2020 in Ecuador.

| Variable | Category | Estimate | SE | OR (95% CI) | Signif. |
| --- | --- | --- | --- | --- | --- |
| Intercept |  | -4.175 | 0.356 |  |  |
| ‡ Swill feeding | No | - | - | 1 |  |
|  | Yes | 2.228 | 0.305 | 9.28 (5.30–17.68) | *** |
| ‡ Notification time | 1–7 days | - | - | 1 |  |
|  | >7 days | 0.778 | 0.147 | 2.18 (1.63–2.90) | *** |
| ‡ Animal entry (last 30 days) | No | - | - | 1 |  |
|  | Yes | 0.734 | 0.149 | 2.08 (1.55–2.79) | *** |
| ‡ Vaccination record CSF | Yes | - | - | 1 |  |
|  | No | 0.629 | 0.139 | 1.88 (1.43–2.47) | *** |
| † Natural Region | Highlands | - | - | 1 |  |
|  | Coastal | 0.627 | 0.169 | 1.87 (1.34–2.61) | *** |
|  | Amazon | -0.127 | 0.169 | 0.88 (0.63–1.23) |  |
| § Age (months) | 1–3 | - | - | 1 |  |
|  | 3–6 | 0.460 | 0.158 | 1.58 (1.16–2.16) | ** |
|  | >= 7 | 0.340 | 0.206 | 1.40 (0.93–2.09) |  |

Chi-sqr: 0.0128 \*, GOF Hosmer-Lemeshow: 0.788, AUC: 0.746, D2: 0.148, R2: 0.143. Significance: \*\*\* 0.001, \*\* 0.01. Levels of risk characterisation: † Population level, ‡ Herd level, § Animal level.

Factors that substantially increased the odds of CSF occurrence at the herd level were swill feeding (OR 9.28), notification time (OR 2.18), entry of animals in the last 30 days (OR 2.08) and lack of CSF vaccination (OR 1.88). At the population level: being in a coastal region (OR 1.87) and at the animal level: age of animals between 3–6 months (OR 1.58) (Table 4, Figure 4). The final logistic model presented good fit (GOF=0.79, AUC=0.75). Individual collinearity diagnostics for each variable resulted in individual GVIFs below 1.062. There was no outlier with a significant influence on model fitting, according to the Bonferronni outlier test (p=0.0072); also, there was no correlation between residuals.

**Figure 4.**
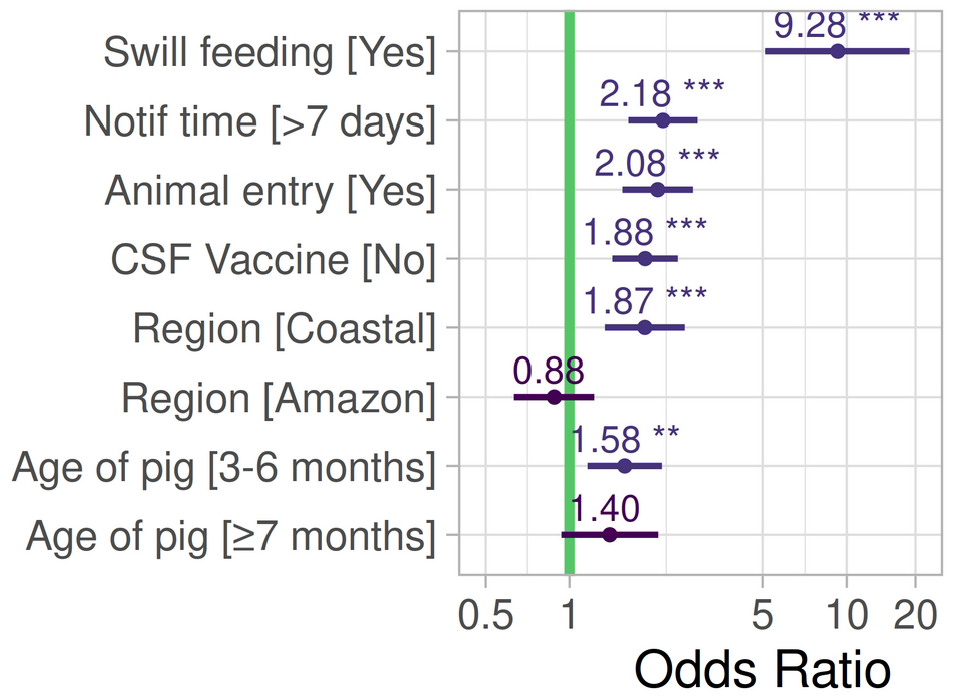
Variables associated with the odds of classical swine fever in Ecuador 2014-2020.

### 4.4 Spatiotemporal descriptive analysis

The Ecuadorian administrative division has 1,040 parishes; 16 were removed because of being islands and 18 were located in the Amazon rainforest with no record of domestic pigs. The length of the neighbouring areas was 4,024 (1,006 for each year), with 271 (6.49%) imputed gaps. When comparing the imputed data with the original dataset, they were not significantly different (t-test: p=0.55). The final neighbour list contained 1,006 parishes with an average of 5.71 parish neighbours. The average parish area was 198.03 ± 249.48 km^2^, with a range from 2.23 to 2,429.64 km^2^. The annual average of registered premises was 115,411.8, housing an average of 1,633,922 pigs.

The annual average of CSF vaccine doses between 2017 and 2019 was 2,375,290, expressed as vaccine doses per square kilometre from 15.7 in 2017 to 23.0 in 2020; the highest average vaccination coverage was 81% in 2019, and the lowest was 60% in 2017. The number of applied doses increased from 1.8 millions in 2017 to 2.4 millions in 2020. The annual average of premises was 115,411 (Table 5).

**Table. 5.** Centrality measures of model variables (fixed effects) aggregated by parish distribution in Ecuador.

| Variable | 2017 |  | 2018 |  | 2019 |  | 2020 |  |
| --- | --- | --- | --- | --- | --- | --- | --- | --- |
|  | Average | Median (max) | Average | Median (max) | Average | Median (max) | Average | Median (max) |
| Doses CSF/km<sup>2</sup> | 15.74 | 2.31 (976.5) | 23.78 | 4.15 (1057.6) | 27.63 | 5.08 (1,282) | 23.0 | 4.2 (1,477.3) |
| Population of pigs | 1,671.3 | 284 (224,448) | 1,948.7 | 391 (256,107) | 2,030.4 | 447 (254,042) | 1,867.1 | 434.5 (227,186) |
| Vaccine coverage % | 60 | 64 (129) | 71 | 80 (108) | 81 | 100 (105) | 70 | 73 (103) |

What stands out in figure 5 is the decline of the number of observed cases, corresponding to 39 in 2017, 60 in 2018, 33 in 2019 and 13 in 2020; it is possible to observe a significant reduction in the number of cases specially in the highlands over the years.

**Figure 5.**
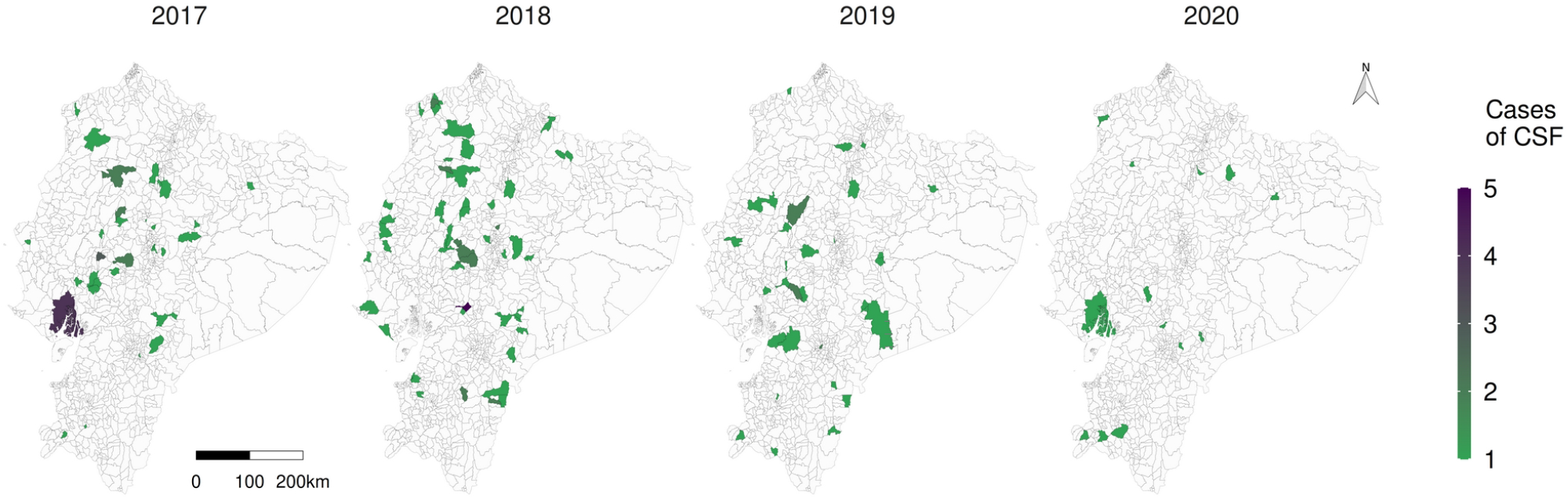
Representation of the number of observed CSF cases (number of positive premises) in Ecuador grouped by parish from 2017 to 2020.

Figure 6 reveals that there was a marked higher density on the number of doses of CSF applied by square kilometre in the western centre (Santo Domingo), the north (Carchi), west south (El Oro) and the central highlands (Cotopaxi, Chimborazo), this general pattern repeated over the years, however there were 105, 78, 52 and 62 parishes without vaccination coverage in 2017, 2018, 2019 and 2020 (white on Figure 6).

**Figure 6.**
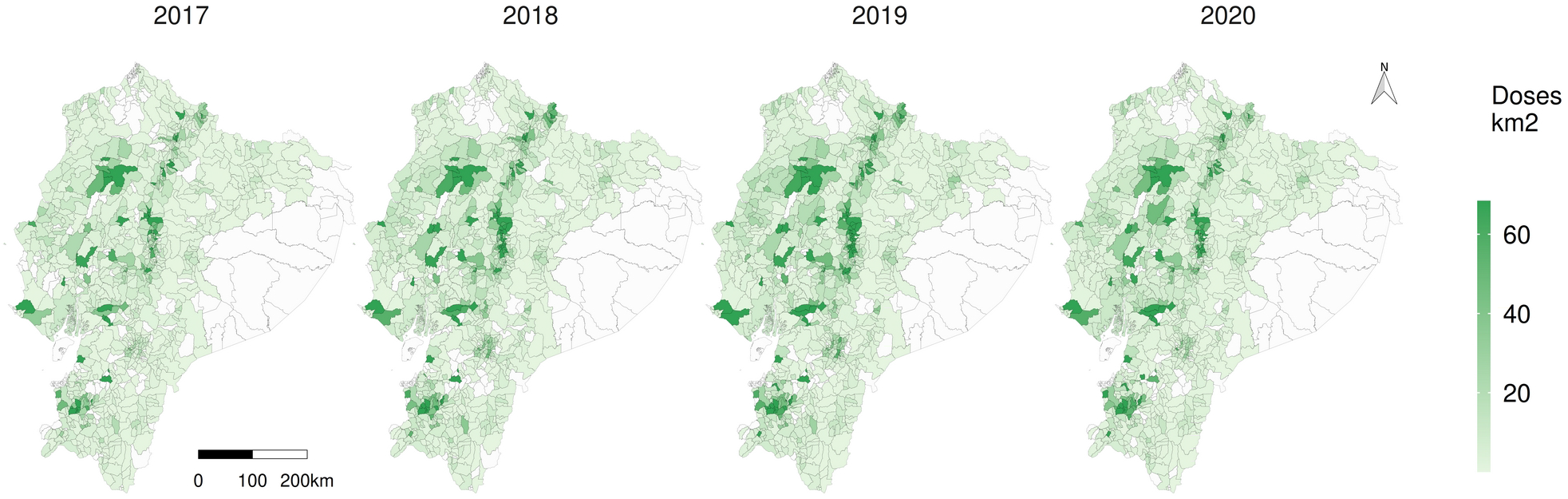
Classical swine fever vaccination density per square kilometre in Ecuador. Polygons of the parishes on the map. White parish without applied doses.

Temperature and precipitation in Ecuador are modulated by the Andes mountains, warmer temperatures on the Amazon and Coastal regions and cooler temperatures on the highlands. There is a range difference of 11 °C in the Coastal, 20.5 °C in the Highlands and 18.9 °C in the Amazon. Precipitation on the Amazon is almost 3 times higher than the highlands and two times compared with the Coastal (Table 6).

**Table 6.** Descriptive measures of covariants spatiotemporal model in Ecuador.

| Covariate | Coastal | Highlands | Amazon |
| --- | --- | --- | --- |
| | Mean ± SD (range) | Mean ± SD (range) | Mean ± SD (range) |
| Doses vac. Km<sup>2</sup> | 18.93 ± 69.83 (0–636.9) | 19 ± 61.01 (0–976.5) | 1.77 ± 2.99 (0–15.26) |
| Temperature | 24.36 ± 1.63 (15–26) | 14.26 ± 4.43 (4.6–25.1) | 20.60 ± 4.19 (6.8–25.7) |
| Precipitation | 1362.24 ± 712.20 (122–3253) | 1012.89 ± 434.16 (432–3824) | 2718.53 ± 957.47 (722–4482) |

### 4.5 Spatiotemporal Relative Risk

The annual average relative risk dropped from 4.01 in 2017 to 1.30 in 2020. Regarding the doses of vaccine applied per km^2^, they behaved as an expected protective factor, which means that an increase in one SD in the doses applied per kilometre decreased the risk by 33%. Temperature was a risk factor, considering that an increase in one SD in temperature degree increased the risk by 16.7%. Precipitation had no effect: RR=1.00 (1.00–1.001) (Table 7), and the spatial distribution of risk is shown in Figure 8.

**Table 7.**
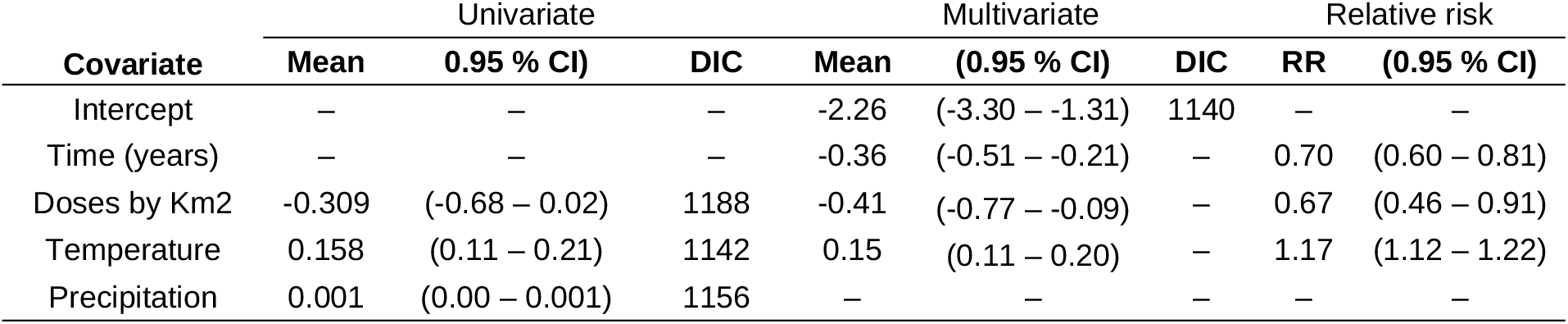
Summary statistics of the effect of covariates on the estimated risk (RR) of CSF in a spatiotemporal Bayesian model.

**Figure 7.**
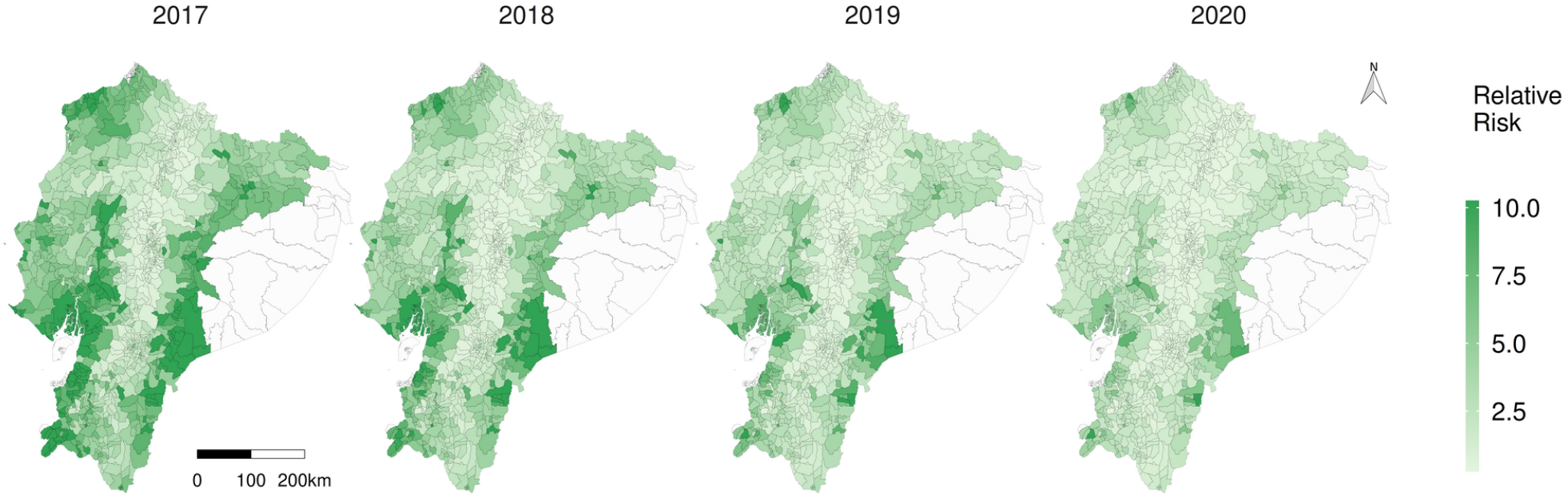
Spatiotemporal representation of the relative risk (RR) of CSF in Ecuador.

**Figure 8.**
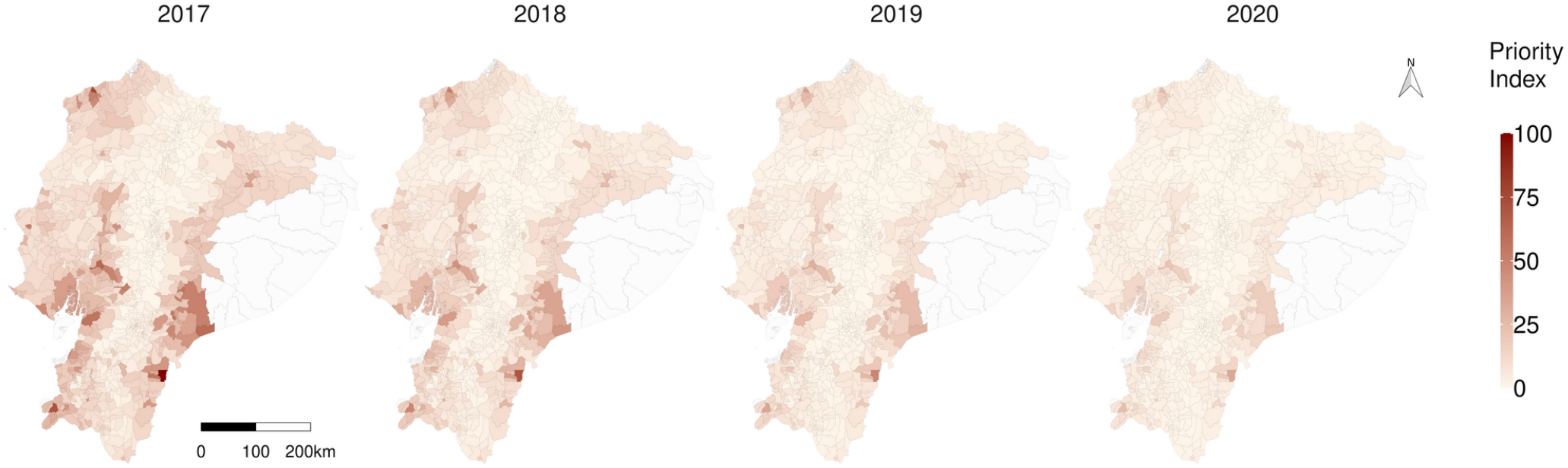
Spatiotemporal representation of the priority index (PI) to fight classical swine fever in Ecuador.

The proportion of variance explained by each component was 57% for the random effect (iid), a major contributor to the explained variance, and 43% for the spatiotemporal (besag). The DIC mean deviance was 1,140 and the effective number of parameters was 136.7.

Hot spots of increased risk were spatially identified on the map, in the coastal southwestern and also the southeastern Amazon, reducing the risk over the years (Figure 7).

According to the priority index (PI), prime concern parishes are located in the eastern Amazon as *Tundayme*, followed by *Tachina* in the northwestern and *Paletilla* in the southern zone (Figure 8). The provinces with higher risk, considering the average RR per province in the year 2020, were Morona Santiago (3.68), Los Rios (3.12) and Santa Elena, (3.07).

## 5. Discussion

When countries mount resource-intensive control strategies for high-impact disease as CSF but fail to reach the goal of control and elimination, a deeper analysis of the disease dynamics and the implemented control interventions is needed to identify strategic intervention points. Despite the fact that in general terms many CSF risk factors are known, their relevance in the specific setting of a pig sector and the respective control program is ideally assessed using all available data.

Swill feeding is one of the main risk factors for CSF transmission; it is common and rooted in the cultural tradition of backyard producers (43); therefore, it is very likely to be a key disease driver of the disease in endemic areas of Andean countries. The Agricultural Health Law of 2019 (44), established best practices for animal feed, but lacked specific regulations on swill. Consideration needs to be given to promoting risk-reducing practices such as heat treatment (45) and stricter regulations that prohibit the use of animal protein as a feed source for pigs.

Vaccination misreporting could be related to the producer’s lack of knowledge regarding veterinary treatments linked with injections (vaccination, iron supplementation in piglets, deworming or other). Fear associated with owners’ legal responsibilities and misunderstandings during the interviews may also lead to misreporting (46). In Indonesia, vaccination against CSF resulted in an increased risk of CSF due to inaccurate vaccination claims (23); considering these facts, reporting behaviour could be further analysed as an early target of the surveillance programme (47); suggesting that communication and health education activities might be advisable to improve producers’ understanding of animal disease prevention and control practices.

Our findings on increased risk at the age of 3-6 months could be related to the fact that young animals might be more likely to be exposed to CSFV because this is the age at which they are normally marketed. The age with increased risk for CSF also reflects a complicated age from the immunological perspective. Maternal immunity fades out after three months (48), and animals not vaccinated become susceptible just as animals vaccinated too early where maternal antibodies interfere with the vaccination. Nowadays, the established recommendation for piglets in Ecuador is a primary vaccination at relatively early 45 days and revaccination every 6 months. This practice might have to reassess once targeted sero-surveillance studies to clarify the effects of vaccination ages and herd immune status.

Maternal-derived antibody (MDA) interference is the most common factor affecting the induction of protective immunity against CSFV; in Thailand the vaccination program has been implemented for decades without achieving eradication (49). In addition, emergency vaccination protocols implemented in very young piglets, especially during an outbreak, could be further analysed. It would be necessary to evaluate diagnostic tools (rapid test) (50) that could detect non clinical, persistent CSF forms in the field, as well as apply vaccination serological monitoring tools (51).

Cases occurred on farms that recorded vaccination, this could be related to illegal movements of unvaccinated animals (52); and, they could also be a result of vaccination failures due to poor handling and malpractice as evidenced in Colombia (4,19). The risk associated with temperature in the spatiotemporal analysis, could be related to vaccination failures, as the vaccine cold-chain in regions such as the Amazon and the Coastal with average temperatures above 20 degrees Celsius, which corroborates that vaccination strategies alone are not sufficient to eradicate the disease (53,54); possible antigenic alteration due to vaccination pressure and their effects on disease epidemiology could also be further analysed (55,56).

The identified individual parish risk could help identify neglected territories; as in many developing countries with limited resources for disease control, prioritisation is often done on the basis of historic surveillance information; therefore, reduced surveillance sensitivity may leave areas of high risk unnoticed. Our model included all parishes and considered the influence of their neighbours to improve the predictions (57). Concepts such as spatial RR or excess risk might be difficult to interpret outside the scientific community, but the priority index (PI) could facilitate understanding and communication of which parishes should be prioritised.

The identification of the risk factors should respond to the initial demand of the NVS and contribute to the implementation of a risk-based surveillance strategy for CSF. As risk factors are specific for each disease new studies could be implemented using depurated data and methodology for prevalent diseases where symptomatology could be confused with CSF such as the porcine reproductive and respiratory syndrome (58,59), also prepare the surveillance system for re-emerging diseases such as African swine fever, currently detected in Central America (60,61).

In total, the results indicate once again the complexity a CSF control program is facing, particularly if the pig sector is diverse and comprises a large share of farms falling under the subsistence or backyard category. Here, NVS faces risky production methods combined with reduced knowledge on disease prevention and compliance with sanitary regulations.

## Author contributions

A. Acosta, F. Ferreira and K. Depner conceived the study; C. Imbacuan and L. Burbano coordinated the data collection; K. Dietze, G. Osowski and A. Acosta curated the data and wrote the manuscript; O. Baquero participated in the spatiotemporal analysis; A. Acosta conducted data processing and coding; All authors discussed the results and critically reviewed the final manuscript.

## Acknowledgements

The authors would like to thank Dr. Javier Vargas for his leadership in the implementation of the national swine health project, Dr. Stalin Vasquez for internal coordination and the National Veterinary Service for their collaboration and data provision. The views expressed are those of the author(s) and not necessarily of the Veterinary service or the Ecuadorian Ministry of Agriculture.

## Data availability statement

The surveillance data contain private information of each Ecuadorian pig farmer and therefore cannot be made available due to legal restrictions.

## Ethics statement

The authors confirm that the ethical policies of the journal, as noted on the journal’s author guidelines page, have been adhered to. No ethical approval was needed as this work did not involve animal sampling or questionnaire data collection by the researchers.

